# Genetic structure and diversity of *rfb* locus of pathogenic species of genus *Leptospira*

**DOI:** 10.1101/2023.03.23.533916

**Authors:** L. C. A. Ferreira, L. F. A. Ferreira Filho, M. R. V. Cosate, T. Sakamoto

## Abstract

Considered a globally important zoonotic bacterial disease, leptospirosis affects both humans and commercially important animals. It is transmitted through direct or indirect exposure to the urine of infected animals and is a major public health challenge in regions with heavy rainfall, floods, and poor socioeconomic conditions. The genus *Leptospira* has 67 species, which can be grouped into pathogenic and saprophytic groups. Serological classification based on antigenic characteristics is important in epidemiology and clinical analysis but is laborious, requires infrastructure and specialized labor, and takes days to obtain results. In this study, we aimed to find genetic patterns associated with the serological classification of *Leptospira* to propose molecular markers for classifying *Leptospira* samples at the serogroup level. For this, we used genomic data of 722 samples distributed in 67 species in public databases and compared the gene composition of their *rfb* locus. Clustering analysis was able to group samples into five major classes that share similarities in both the serological and genetic composition of the *rfb* locus. We also identified some syntenic blocks in the internal region of the *rfb* locus and patterns of presence and absence of these blocks which can be used to determine the serogroup of a sample. Our findings can assist the development of molecular strategies for the serological identification of *Leptospira* samples, which could be more rapid and accurate than the current method.

## Introduction

Leptospirosis is a zoonotic bacterial infectious disease that affects both humans and commercially important animals. The disease is widely distributed geographically and transmitted through direct or indirect exposure to the urine of infected animals, representing a major public health challenge in regions with heavy rainfall, floods, and poor socioeconomic conditions^1,2^. The causative agent of the disease is the pathogenic species of the genus *Leptospira*, a highly motile gram-negative bacterium thanks to its periplasmic flagella, and which is found in water and soil worldwide^3,4,5^. According to the Centers for Disease Control and Prevention^6^, the disease is responsible for over one million cases annually worldwide and, since it is considered a neglected zoonosis^7^, the actual numbers are potentially even higher.

Until 1989, *Leptospiras* were divided into only two species, *L. interrogans* and *L. biflexa*, with pathogenic strains represented exclusively by the species *L. interrogans* and saprophytes by *L. biflexa*^8^. Later, with the advances in diagnostic methods, *Leptospira* samples has been classified using two distinct approaches : the serological classification, which is based on their antigenic characteristics called serovar (srv), and taxonomic classification, in which we could presume the evolutionary relationship between samples and is based on the DNA sequence similarity. It is interesting to note that the two forms of classification do not correspond to each other and each one is used for different purposes. Serological classification is important for epidemiological studies of disease and vaccine development, while taxonomic classification is important in evolutionary studies of the genus9,10,11.

Recent studies on the evolution and taxonomic classification of genus *Leptospira* have classified the 68 species identified so far into two major clades in which one of them comprises all pathogenic species, and the other, all saprophytic species. These clades are further subdivided into two subclades each. The pathogenic clade is subdivided into P1 and P2 subclades, which are also known as pathogenic and intermediate group, respectively., and thesaprophytic clade is subdivided into S1 and S2^12^.

*Leptospira* samples can also be classified into more than 30 serogroups and 300 serovars^13^. The serological classification is based on the structure of the lipopolysaccharides (LPS). LPS is a complex molecule present in the outer layer of the outer membrane of bacteria such as *Leptospira*, constituting the main immunodominant antigens of these organisms^14^. Differences in sugar composition and orientation in the LPS orientation are the main characteristics that distinguish different serovars^15^. The genomic region responsible for LPS antigen synthesis is the *rfb* locus^16^, and the genes in this region are the main genetic factors associated with the serological classification of the genus *Leptospira*^17^.

Leptospirosis is a disease that is often underdiagnosed due to its multiform and nonspecific presentation, and it is very common for leptospirosis to be confused with other diseases that have febrile states such as malaria and hepatitis, especially viral hemorrhagic fevers such as dengue^18^. Therefore, it is essential that laboratories have the ability to accurately classify the serology of pathogenic strains of the *Leptospira* genus in order to direct the most appropriate treatment and prophylactic measures through vaccines. However, the methods used to perform the serological classification, such as Microscopic Agglutination Test (MAT), are time-consuming and expensive.

Therefore, there is an urge to develop more effective methods that could perform serological identification.Molecular-based methods, such as PCR analysis and DNA sequencing, are promising alternatives to the current methods. Although there are suggestions in the literature, there are still no reliable methods for performing this procedure. This is due to the still scarce knowledge about the genetic basis associated with serological classification.

The number of genomes from the genus *Leptospira* has been significantly increasing in public databases over the years, as a result of the increased processing capacity of sequencing technologies^20^. All of this available data allows us to conduct studies that aim to find the genetic basis associated with the serological classification of *Leptospira spp*. strains. Therefore, the aim of this study was to further understand the genetic basis underlying the serological classification in *Leptospira* to aid the development of molecular strategies for serological identification, For this we accessed the genomic data of more than 700 samples of *Leptospira* and performed a comparative analysis of the structure and organization of their *rfb* locus.

## Results

### Hierarchical clustering of *Leptospira* samples based on the genetic composition of the *rfb* locus

The *rfb* locus of the genus *Leptospira* comprises a broad variety of genes. To compare the gene composition of this locus among the samples used in this study, we performed a hierarchical clustering analysis using a presence-absence table of *rfb* locus genes. The table was created based on the ortholog clusters inferred from the proteomes of 722 *Leptospira* samples. This generated a total of 14,067 orthogroups, of which 395 contained genes that are part of the *rfb* locus. After submitting the presence-absence table to the clustering method, a heatmap and 2-dimensional hierarchical clustering were generated (Fig. 1).

**Figure 1:**
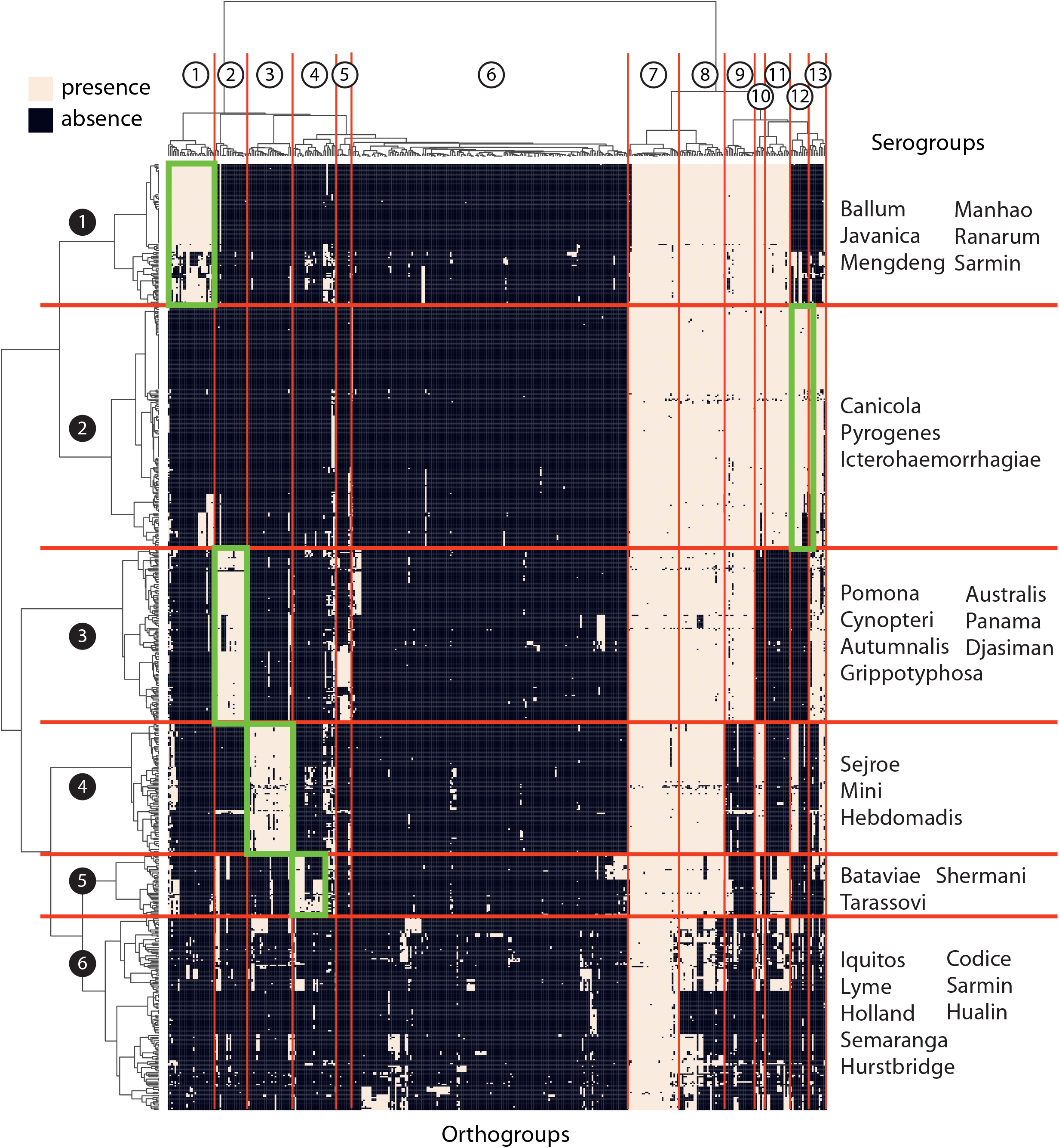
Hierarchical clustering of *Leptospira* samples based on the genetic composition of the *rfb* locus. Values are represented on a binary scale, with darker color indicating zero (absence) and lighter color indicating a value of 1 (presence). On the x-axis, the tree illustrates how the orthogroups are grouped, while on the y-axis, the tree represents the clustering of the samples. The samples were subdivided into six (sample) clusters and orthogroups were subdivided into 13 (ortho)clusters. Areas highlighted in green represent orthogroups specific to a sample cluster.

In this analysis, samples were grouped into six well-defined clusters (cluster 1 to 6), which will be referred to here as sample clusters. The distribution of serogroups among the sample clusters shows that samples belonging to one serogroup are, in general, in the same cluster (Table 1). Most of the samples from the serogroups Icterohaemorrhagiae (n=117),

**Table 1:** Distribution of serogroups in the analyzed samples, highlighting the clustering of samples belonging to the same serogroup.

| Serogroups | Clusters |  |  |  |  |  | Total |
| --- | --- | --- | --- | --- | --- | --- | --- |
|  | 1 | 2 | 3 | 4 | 5 | 6 |  |
| Ballum | 66 |  |  |  |  |  | 66 |
| Javanica | 9 |  |  |  |  |  | 9 |
| Manhao | 3 |  |  |  |  |  | 3 |
| Mengdeng | 3 |  |  |  |  |  | 3 |
| Ranarum | 2 |  |  |  |  |  | 2 |
| Sarmin | 1 |  |  |  |  |  | 1 |
| Canicola | 1 | 9 | 1 |  |  |  | 11 |
| Icterohaemorrhagiae |  | 116 | 1 |  |  |  | 117 |
| Pyrogenes |  | 20 | 1 |  |  |  | 21 |
| Australis |  |  | 9 |  |  |  | 9 |
| Autumnalis | 2 |  | 9 |  |  |  | 11 |
| Cynopteri |  |  | 1 |  |  |  | 1 |
| Djasiman |  |  | 1 |  |  |  | 1 |
| Grippotyphosa |  |  | 24 |  |  | 1 | 25 |
| Panama |  |  | 2 |  |  |  | 2 |
| Pomona |  |  | 27 |  |  |  | 27 |
| Hebdomadis |  |  |  | 3 |  |  | 3 |
| Mini |  |  |  | 6 |  |  | 6 |
| Sejroe |  |  |  | 42 |  |  | 42 |
| Bataviae |  |  |  | 1 | 11 |  | 12 |
| Shermani |  |  |  |  | 2 |  | 2 |
| Tarassovi |  |  |  |  | 10 |  | 10 |
| Codice |  |  |  |  |  | 1 | 1 |
| Holland |  |  |  |  |  | 1 | 1 |
| Hualin |  |  |  |  |  | 1 | 1 |
| Hurstbridge |  |  |  |  |  | 2 | 2 |
| Iquitos |  |  |  |  |  | 2 | 2 |
| Lyme |  |  |  |  |  | 2 | 2 |
| Sarmin |  |  |  |  |  | 1 | 1 |
| Semarang |  |  |  |  |  | 4 | 4 |
| Unknown | 23 | 38 | 56 | 49 | 42 | 116 | 324 |
| Total | 110 | 183 | 132 | 101 | 65 | 131 | 722 |

Ballum (n=66), and Sejroe (n=46), which are the most frequent serogroups in the analysis, were grouped into the sample clusters 1, 2, and 4, respectively. It is noteworthy that samples belonging to different species could belong to the same serogroups (Table 2). For instance, we could find five different species among samples belonging to the serogroup Mini, *L. interrogans, L. kirschneri, L. borgpetersenii, L. santarosai*, and *L. mayottensis*, and all of them were in the same cluster. We also verified the distribution of species in the six clusters (Table 3) and found that five of them consist only of samples from the P1 clade. Cluster 6, which is the most heterogeneous group with respect to species, contains samples from all four *Leptospira* clades (P1, P2, S1, S2). Among the samples of the P1 clade, we could note that there is no clear pattern of their distribution among the clusters. The samples of species *L. interrogans* (n=314), *L. borgpetersenii* (n=130), and *L. santarosai* (n=40), which are the most frequent species, can be found in five, four, and five clusters, respectively. All these observations indicate that the clustering generated in this work is independent of taxonomic classification and strongly associated with serological classification.

**Table 2:** Distribution of serogroups among the pathogenic species of *Leptospira*

| serogroups | *Lint* | *Lnog* | *Lale* | *Lkir* | *Lbor* | *Lsan* | *Lwei* | *Lmay* | *Lals* | *Lkme* |
| --- | --- | --- | --- | --- | --- | --- | --- | --- | --- | --- |
| Australis | 8 | 1 |  |  |  |  |  |  |  |  |
| Autumnalis | 4 | 2 | 2 | 3 |  |  |  |  |  |  |
| Ballum |  |  |  |  | 65 |  |  |  |  |  |
| Bataviae | 11 | 1 |  |  |  |  |  |  |  |  |
| Canicola | 11 |  |  |  |  |  |  |  |  |  |
| Cynopteri |  |  |  | 1 |  |  |  |  |  |  |
| Djasiman | 1 |  |  |  |  |  |  |  |  |  |
| Grippotyphosa | 11 |  |  | 10 |  | 4 |  |  |  |  |
| Hebdomadis | 1 |  |  |  |  |  | 2 |  |  |  |
| Icterohaemorrhagiae | 117 |  |  | 1 |  |  |  |  |  |  |
| Javanica |  |  |  |  | 6 | 3 |  |  |  |  |
| Manhao |  |  | 2 |  |  |  | 1 |  |  |  |
| Mengdeng |  |  |  |  |  |  | 3 |  |  |  |
| Mini | 1 |  |  | 2 | 1 | 1 |  | 1 |  |  |
| Panama |  | 2 |  |  |  |  |  |  |  |  |
| Pomona | 19 |  |  | 8 |  |  |  |  |  |  |
| Pyrogenes | 21 |  |  |  |  |  |  |  |  |  |
| Ranarum |  |  |  |  |  |  | 1 |  | 1 |  |
| Sarmin |  |  | 1 |  |  | 1 |  |  |  |  |
| Sejroe | 8 |  |  |  | 33 | 1 |  |  |  |  |
| Shermani |  |  |  |  |  | 2 |  |  |  |  |
| Tarassovi |  |  | 1 |  | 7 | 1 | 1 |  |  | 1 |

**Table 3:** Distribution of *Leptospira* species in each cluser.

| Clades | Species | Clusters |  |  |  |  |  | Total |
| --- | --- | --- | --- | --- | --- | --- | --- | --- |
|  |  | 1 | 2 | 3 | 4 | 5 | 6 |  |
| P1 | *L. adleri* |  |  |  |  |  | 3 | 3 |
|  | *L. alexanderi* |  | 1 |  | 5 |  |  | 6 |
|  | *L. alstonii* |  |  |  | 4 | 1 |  | 5 |
|  | *L. barantonii* |  | 2 |  |  |  |  | 2 |
|  | *L. borgpetersenii* |  | 11 | 40 | 77 |  | 2 | 130 |
|  | *L. dzianensis* |  | 2 |  |  |  | 4 | 6 |
|  | *L. ellisii* |  |  |  |  |  | 2 | 2 |
|  | *L. gomenensis* |  |  |  |  |  | 4 | 4 |
|  | *L. interrogans* | 178 | 15 | 29 | 4 | 88 |  | 314 |
|  | *L. kirschneri* | 4 | 1 | 5 |  | 25 |  | 35 |
|  | *L. kirschneri* |  | 7 |  |  |  |  | 7 |
|  | *L. mayottensis* |  | 1 | 3 | 2 |  |  | 6 |
|  | *L. noguchii* |  | 5 |  | 1 | 11 |  | 17 |
|  | *L. putramalaysiae* |  | 2 |  |  |  |  | 2 |
|  | *L. santarosai* |  | 11 | 15 | 5 | 7 | 2 | 40 |
|  | *L. stimsonii* |  | 2 |  |  |  |  | 2 |
|  | *L. tipperaryensis* |  | 1 |  |  |  |  | 1 |
|  | *L. weilii* |  | 1 | 8 | 11 |  |  | 20 |
|  | *L. yasudae* |  | 1 |  | 0 |  | 1 | 2 |
|  | Total | 182 | 63 | 100 | 109 | 132 | 18 | 604 |
| P2 |  |  |  |  |  |  | 40 | 40 |
| S1 |  |  |  |  |  |  | 67 | 67 |
| S2 |  |  |  |  |  |  | 5 | 5 |
|  | *Leptospira sp.* | 1 | 2 | 1 | 1 |  | 1 | 6 |

The analysis also involved genomic data from samples that did not have serological identification, which totalized 324 samples. Despite the lack of serological data, the inclusion of these samples is interesting to verify other organizations and structures of the *rfb* locus. All these samples were grouped in one of the 6 clusters previously mentioned, suggesting that they could share antigenic similarity with serologically characterized samples and could be classified into one of the existing serogroups. It is of high interest that serological identification of these samples is performed to confirm the findings of this study.

Hierarchical clustering analysis also allowed us to identify the orthogroups that are characteristic of each cluster (Fig. 1). To do this, we established a threshold on the dendogram that separated the orthologous groups into 13 clusters, which will be referred to here as orthoclusters. In this analysis, it was possible to identify the existence of at least five orthoclusters (1, 2, 3, 4, and 12) that characterize a sample cluster (highlighted in green in Fig. 1).

### Comparative analysis of structure and organization of *Leptospira rfb* locus

To visualize the gene organization and to identify patterns among different profiles of the *rfb* locus in the genus *Leptospira*, we generated diagrams illustrating the genetic structure and organization of the *rfb* locus in representatives of each serogroup in the analysis (Fig. 2). In these diagrams, we could verify regions that are conserved in most samples of the serogroups represented and regions that vary considerably in the gene composition. The 3’-terminal region of the samples from clusters 1 to 5 is one of the conserved regions. In addition to several genes that encode sugar-modifying enzymes, such as methyltransferases and glycosyltransferases, this region comprises genes responsible for the O-antigen processing and synthesis through the Wzy-dependent pathway, and for the dTDP-rhamnose biosynthesis (*rfbC, rfbD, rfbB, rfbA*), which are involved in the assembly of LPS. Genes found in all representative samples are the MarR, which is a transcriptional regulator that initiates the *rfb* locus, and the GDP-L-fucose synthase. In the 5’ region of the *rfb* locus, we could also verify a set of genes encoding sugar-modifying enzymes conserved in samples from the clusters 1, 2, and 3.

**Figure 2:**
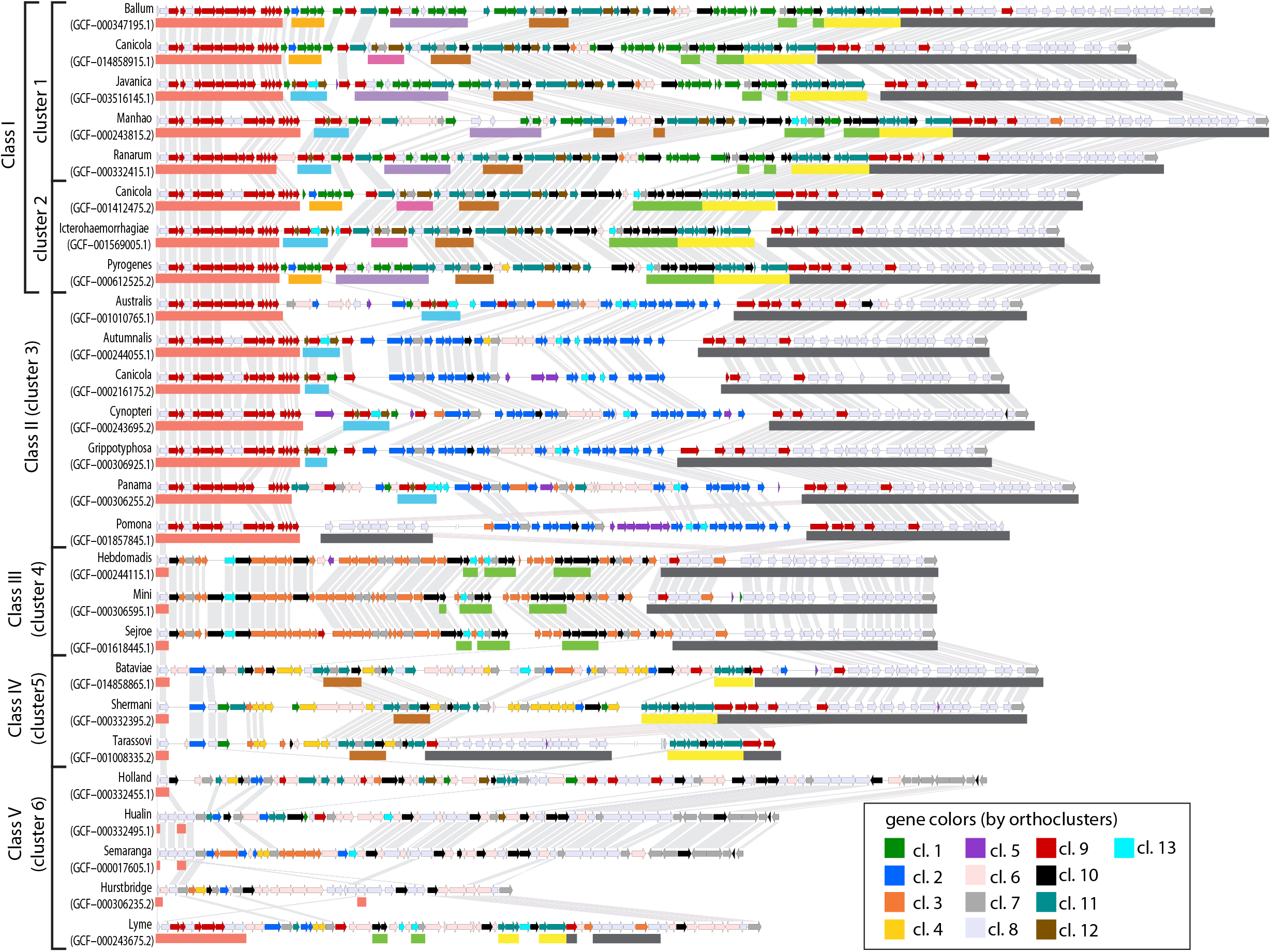
Genetic organization of the *rfb* locus in representative sample of each serogroup. Genes that have the same color belong to the same cluster. The samples were ordered according to their similarities in *rfb* locus gene composition verified in the hierarchical clustering analysis. Gene pairs from adjacent samples that share more than 50% identity are linked. Each gene is colored according to the orthocluster to which it belongs. Bars below some gene blocks highlight syntenic blocks found in the locus.

The innermost region of the *rfb* locus has a more diverse gene composition among the analyzed samples. This region consists mainly of genes encoding enzymes that add or modify sugar during the O-antigen biosynthesis. Comparing the gene composition of the locus between samples from different clusters, we can see that each cluster presents distinctive gene composition, except for the samples from clusters 1 and 2. For this reason, we decided to join these two clusters and named the new group Class I. Clusters 3, 4, 5, and 6 were renamed respectively as class II, III, IV, and V.

Interestingly, there are several small syntenic blocks that may be conserved in samples from different species, classes, and serogroups (Fig. 2). The different combinations of these syntenic blocks at the *rfb* locus suggest that they are determinants in expressing the serogroup of a sample. Taking the Icterohaemorrhagiae serogroup (Class I) sample, it can be seen that there is a set of genes (highlighted with light blue bar, Fig. 2) that can differentiate samples of this serogroup from Canicola and Pyrogenes. But since this same set of genes can be found in other samples, such as in the serogroup Javanica (Class I), Autumnalis, Cynopteri, and Panama (Class II), a second set of markers is needed. This may be a gene from a set that is found only in samples from serogroup Icterohaemorrhagiae and Canicola (highlighted with pink bar, Fig. 2). The specificity of these gene sets in serogroup Icterohaemorrhagiae was confirmed by verifying their conservation among the Icterohaemorrhagieae samples and their absence in samples from other serogroups (Fig. 3). Genes found in these blocks are promising molecular markers for serogroup identification. The occurrence of these syntenic blocks also suggests that this region is prone to lateral transfer events. Phylogenetic analysis is of great interest to understand the events of change or the emergence of new serogroups.

**Figure 3:**
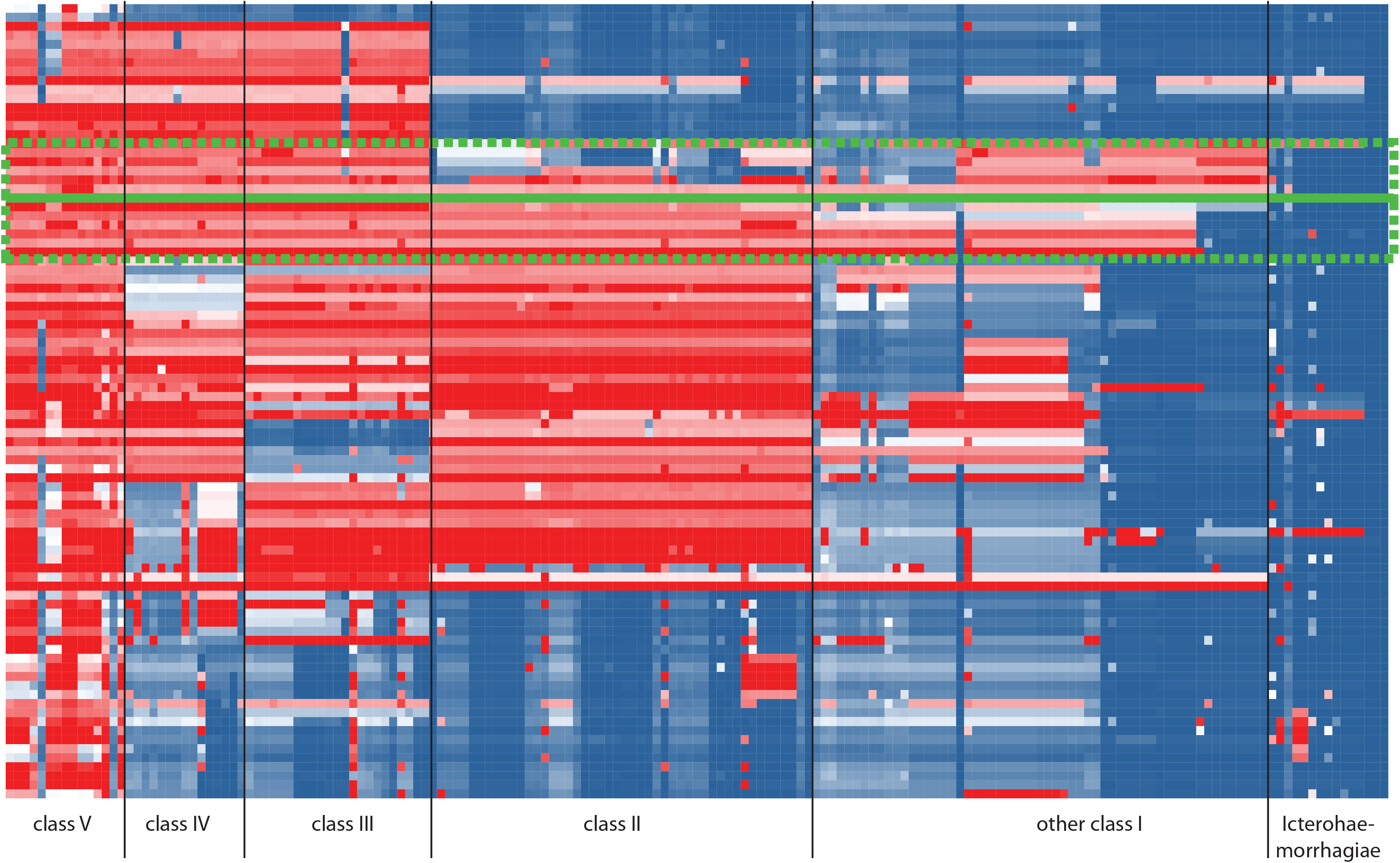
Analysis of sequence similarity of 88 proteins that make up the *rfb* locus of *L. interrogans* (srg Icterohaemorrhagiae) with other Leptospira strains.

Although the genes involved in the biosynthesis of the O-antigen usually cluster at the *rfb* locus, some samples presented segments of this locus in other genome regions. Among them are the samples *L. interrogans* srg Pomona srv Pomona str AKRFB and *L. santarosai* srg Tarassovi str U164 (Fig. 2). In both samples, we could notice long segments containing genes homologous to the internal genes of the *rfb* locus of other samples, but located outside the locus. One could suggest that we are dealing with samples that have low sequencing quality and, consequently, with assembly errors. However, there are samples with well-assembled genomes, that also showed this characteristic. This observation should be considered when attempting to identify the serology of samples using genomic data.

The analyses performed in this work also suggest revisions in the serological classification of some samples. The *rfb* locus profile of samples *L. alexanderi* srv Erinaceiauriti str 56159 and *L. alexanderi* srv Nanla str 56650, both currently classified in the Autumnalis srg, have a high similarity with the profiles of samples from the Manhao srg (class I), suggesting that the most appropriate classification for them is the Manhao serogroup. In a similar way, the revision in serological classification is also suggested for samples *L. interrogans* str UT234, originally from srg Bataviae into srg Sejroe, and *L. interrogans* str 782, originally from srg Canicola, into srg Grippotyphosa.

## Discussion

Infections caused by pathogenic *Leptospira* often manifest asymptomatically or with non-specific symptoms that can be confused with flu-like syndromes. However, when the disease progresses to a severe form, complications can be life-threatening^21^. The methods available to diagnose leptospirosis are serological tests, such as Microscopic Agglutination Test (MAT) and the IgM detection, and molecular tests, such as pulsed-field gel electrophoresis^24^, variable number of tandem repeat (VNTR) detection^25,26^, restriction fragment length polymorphism (RFLP)^27^, and transposable element quantification^28,29^, and PCR. MAT is considered to be the gold standard since it has high sensitivity and is currently the only method that can reliably identify the antigenic group (serogroup) to which a sample belongs. However, performing MAT is often difficult and requires specialized professionals and infrastructure, since it requires the maintenance of living strains of *Leptospira*. Most developing countries, which are the prevalent area of the disease, do not have an adequate infrastructure to perform the test^30^.

The limitation of using PCR-based methods for serological identification was the lack of knowledge about the genetic basis that defines the serology of a *Leptospira* sample. Currently, with the availability of a large volume of genomic data from different species and serogroups of the genus, this limitation is being surpassed. It is known that genes from *rfb* are the main genes associated with the serology of a sample. This was firstly verified by comparing this locus among samples of *L. interrogans* serovar Hardjo, *L. borgpetersenii* serovar Hardjobovis, and *L. interrogans* serovar Copenhageni^17,31^. In these studies, it was observed that the genes of samples belonging to the same serological class but not to the same species shared high similarity when comparing the genes that comprise this locus. Other evidence of the association of this locus with serological classification was found in subsequent studies, such as the sharing of genes in this locus among samples of the Hurstbridge serovar of the *L. fainei* and *L. broomi* species^1^ and the similarity of the gene profile among samples of the Sejroe, Hebdomadis, and Mini serogroups, which have antigenic affinity^4^. A study involving the main serogroups also verified the similarity of the gene profile of the *rfb* locus among samples classified in the same serogroup^12^.

In this study, we were able to further investigate the diversity of gene profiles in the *rfb* locus and the patterns that determine a serogroup type by accessing the genomic data of 722 *Leptospira* samples distributed across 67 species and 30 serogroups. Here we propose five classes of *rfb* locus profiles (class I, II, III, IV, and V) that group samples of serogroups that might share antigenic affinity. This is supported by observing the clustering of some samples from serogroups that were classified indistinctly in the past. The serogroups Sejroe, Mini, and Hebdomadis, which comprise the class III, were once part of a single serogroup called Hebdomadis, which was separated in 1982^32^. Similarly, the serogroups Autumnalis and Djasiman were part of a single serogroup called Autumnalis until 1982^32^ and they are clustered in the class II.

Since we sought to emphasize the pathogenic group, the present study did not analyzed in further detail the genetic composition of the *rfb* locus of class V. A more detailed analysis of this class is in the project’s perspective. It can also be expected that, with the continuous increase of genomic data generated for other *Leptospira* groups, this class will be extended and subdivided into more levels.

It is important to emphasize that genetic analysis alone is not sufficient to fully determine the antigenic relationship between serogroups. Additional studies, such as cross-agglutination and neutralization assays, are needed to evaluate the antigenic relationship between different serogroups and to determine if the observed genetic similarity reflects significant cross-reactivity in the host immune response. The identification of genetic similarities between different serogroups can be useful for the development of broader and more effective diagnostic techniques and vaccines against leptospirosis. The study’s results also highlight the importance of ongoing efforts to improve the quality of *Leptospira* genomes in order to facilitate the identification of new strains and lineages and improve the effectiveness of existing vaccines and diagnostic methods. Ultimately, the information generated in this study may be useful for understanding the epidemiology and evolution of the studied pathogens, as well as for developing strategies for the diagnosis and prevention of leptospirosis.

## Methods

### Sources of genomic and serological data

The dataset used for this study consisted of a total of 722 proteome data of *Leptospira* samples obtained in May 2021 from the National Center for Biotechnology Information (NCBI) Reference Sequence Database (RefSeq). Data on the serological classification of each sample was obtained from the Bacterial Isolate Genome Sequence Database^33^(http://bigsdb.pasteur.fr/). The samples studied in this work encompass 67 distinct species. Only six samples had taxonomic classification at the genus level. Out of the 722 samples used in the study, 340 samples did not have a determined serogroup classification. The remaining samples were classified into one of 30 serogroups.

We also used the Benchmarking Universal Single-Copy Orthologs (BUSCO) with the spirochaetia_odb10 dataset to measure genome assembly and annotation completeness^34,35^. The minimum and maximum completeness scores found among the samples were 70.7% and 100%, respectively, in which more than 96.5% of samples showed a score above 95%. Despite the existence of a few genomes with low quality, they were not initially discarded since the focus of this study was not to analyze the genome as a whole but rather the rfb locus.

### Ortholog cluster inference and determination of *rfb* locus genes orthogroups

We submitted the entire set of protein sequences of all 722 samples to the Orthofinder^36^ software tool to determine the orthologous groups between the gene sequence sets in the different accesses. The program generated a total of 14,067 orthologous groups, where 34 of them were present in single copy in all samples and 144 were present in all samples, but not necessarily in single copy.

We subsequently identified the orthogroups that compose the *rfb* locus. For this, we selected samples that contained the *rfb* locus assembled in a single contig. This was checked by verifying whether the genes that initiate (MarR) and terminate (DASS) the locus were contained within the same contig. Then, we identified the orthogroups of the genes located between MarR and DASS. We found 200 samples with the intact *rfb* locus from 27 serogroups. Serogroups that were not represented among the 200 samples had the genes belonging to the *rfb* locus extracted through a manual inspection of their genomic data. Of the total of 14,067 orthogroups, 395 contained genes that are part of the *rfb* locus.

### Clustering and visualization of the genetic composition of the *rfb* locus

Hierarchical clustering method was used to group samples with similar genetic profiles in the *rfb* locus. This analysis was performed using the scikit-learn, scipy, and seaborn Python libraries. The ward method was used for clustering and the Euclidean distance metric was used for measuring the distance between samples. LinearDisplay command-line software^37^ was used to generate a diagram of the genetic composition of the *rfb* locus.

The presence and conservedness of a gene were accessed by sequence similarity search using the BLAST algorithm (BlastP)^38^. The identity score between the query and subject proteins was calculated as the product of the percentage of identity and the query coverage of the first high-scoring segment pair (HSP) as performed by Medeiros ^4^.

## References

1. Fouts, D. E. et al. What Makes a Bacterial Species Pathogenic?:Comparative Genomic Analysis of the Genus Leptospira. PLOS Negl. Trop. Dis. 10(2), 1–57. https://doi.org/10.1371/journal.pntd.0004403 (2016).

2. Pinto, G. V. et al. Current methods for the diagnosis of leptospirosis: issues and challenges. J. Microbiol. Methods. 195, 106438. https://doi.org/10.1016/j.mimet.2022.106438 (2022).

3. Ashaiba, A., Arun, A.B., Prasad, K. S., Tellis, R. C.. Leptospiral sphingomyelinase Sph2 as a potential biomarker for diagnosis of leptospirosis. J. Microbiol. Methods. 203, 106621. https://doi.org/10.1016/j.mimet.2022.106621 (2022).

4. Medeiros, E. J. S., Ferreira, L. C. A.; Ortega, J. M., Cosate, M. R. V., Sakamoto, T. Genetic basis underlying the serological affinity of leptospiral serovars from serogroups Sejroe, Mini and Hebdomadis. Infect. Genet. Evol. 103, 1–9. https://doi.org/10.1016/j.meegid.2022.105345 (2022).

5. Costa, F. et al. Global Morbidity and Mortality of Leptospirosis: a systematic review. PLOS Negl. Trop. Dis. 9(9), 1–19. https://doi.org/10.1371/journal.pntd.0003898 (2015).

6. Centers for Disease Control and Prevention (U.S.). Leptospirosis fact sheet for clinicians. Stephen B. Thacker CDC Library collection. https://stacks.cdc.gov/view/cdc/52537 (2018).

7. Clemente, B. M., Pineda-Cortel, M. R., Villaflores, O. Evaluating immunochromatographic test kits for diagnosis of acute human leptospirosis: A systematic review. Heliyon. 8(11), e11829, https://doi.org/10.1016/j.heliyon.2022.e11829 (2022).

8. Levett, P. N. Leptospirosis. Clin. Microbiol. Rev. 14(2), 296–326, https://doi.org/10.1128/CMR.14.2.296-326.2001 (2001).

9. Greene, C. E. Infectious Diseases of the dog and cat. (4. Ed,42 Ch) 431–446 (Athens: Elsevier, 2011).

10. Martins, G. & Lilenbaum, W. The panorama of animal leptospirosis in Rio de Janeiro, Brazil, regarding the seroepidemiology of the infection in tropical regions. BMC Vet. Res. 9(237) https://doi.org/10.1186/1746-6148-9-237 (2013).

11. Ramos, T. M. V., Balassiano, I. T., Silva, T. S. M., Nogueira, J. M. R. Leptospirose: Características da enfermidade em humanos e principais técnicas de diagnóstico laboratorial. Rev. bras. anal. clin. 53(3), 211–218. https://doi.org/10.21877/2448-3877.202102110 (2021).

12. Nieves, C. et al. Horizontal transfer of the rfb cluster in Leptospira is a genetic determinant of serovar identity. Life Sci. Alliance. 6(2), 1–17, https://doi.org/10.26508/lsa.202201480 (2022).

13. Vincent, A. T. et al. Revisiting the taxonomy and evolution of pathogenicity of the genus Leptospira through the prism of genomics. PLOS Negl. Trop. Dis. 13(5),1–25. https://doi.org/10.1371/journal.pntd.0007270 (2019).

14. Collins, P. M. & Ferrier, R. J. Monosaccharides: Their Chemistry and Their Roles in Natural Products. (John Wiley & Sons, 1995).

15. Adler, B. History of Leptospirosis and Leptospira. In: Adler, B. (eds) Leptospira and Leptospirosis. Curr. Top. Microbiol. Immunol. 387. http://doi.org/10.1007/978-3-662-45059-8_1 (Springer, Berlin, Heidelberg, 2015).

16. Mitchison, M. et al. Identification and characterization of the dTDP-rhamnose biosynthesis and transfer genes of the lipopolysaccharide-related rfb locus in Leptospira interrogans serovar Copenhageni. J. Bacteriol. 179(4), 1262–1267. https://doi.org/10.1128/jb.179.4.1262-1267.1997 (1997).

17. La Peña-Moctezuma, A., Bulach, D. M., Kalambaheti, T., Adler, B. Comparative analysis of the LPS biosynthetic loci of the genetic subtypes of serovar Hardjo: Leptospira interrogans subtype Hardjoprajitno and Leptospira borgpetersenii subtype Hardjobovis. FEMS Microbiol. Lett. 177(2), 319–326. https://doi.org/10.1111/j.1574-6968.1999.tb13749.x (1999).

18. Ahmed, N. et al. Multilocus sequence typing method for identification and genotypic classification of pathogenic Leptospira species. Ann. Clin. Microbiol. Antimicrob. 5(28). http://doi.org/10.1186/1476-0711-5-28 (2006).

19. Terpstra, W. J. Human leptospirosis: guidance for diagnosis, surveillance and control. 109p. https://apps.who.int/iris/handle/10665/42667 (World Health Organization, 2003).

20. Buermans, H. P. J. & Dunnen, J. T. Next generation sequencing technology: Advances and applications. Biochim. Biophys. Acta - Mol. Basis Dis. 1842(10), 1932–1941. https://doi.org/10.1016/j.bbadis.2014.06.015 (2014).

21. Maneewatchararangsri, S. et al. Evaluation of a genus-specific rGroEL_1-524_ IgM-ELISA and commercial ELISA kits during the course of leptospirosis in Thailand. Sci. Rep. 11(19785). https://doi.org/10.1038/s41598-021-99377-8 (2021).

22. Faine, Solomon. Leptospira and leptospirosis. (2. ed, 272p) (Medisci, 1999).

23. Haji Hajikolaei, M.R., Rezaei, S., Ghadrdan Mashhadi, A.R., Ghorbanpoor, M. The role of small ruminants in the epidemiology of leptospirosis. Sci. Rep. 12(2148). http://dx.doi.org/10.1038/s41598-022-05767-x (2022).

24. Herrmann, J. L., Bellenger, E., Perolat, P., Baranton, G., Girons, I. S. Pulsed-field gel electrophoresis of NotI digests of leptospiral DNA: a new rapid method of serovar identification. J. Clin. Microbiol. 30 (7), 1696–1702. http://doi.org/10.1128/jcm.30.7.1696-1702.1992 (1992).

25. Salaün, L., Mérien, F., Gurianova, S., Baranton, G., Picardeau, M. Application of Multilocus Variable-Number Tandem-Repeat Analysis for Molecular Typing of the Agent of Leptospirosis. J. Clin. Microbiol. 44(11), 3954–3962. http://doi.org/10.1128/jcm.00336-06 (2006).

26. Majed, Z. et al. Identification of Variable-Number Tandem-Repeat Loci in Leptospira interrogans Sensu Stricto. J. Clin. Microbiol. 43(2), 539–545. http://doi.org/10.1128/jcm.43.2.539-545.2005 (2005).

27. Jung, L. R. C., Bomfim, M. R. Q., Kroon, E. G., Nunes, A. C. Identification of Leptospira serovars by RFLP of the RNA polymerase beta subunit gene (rpoB). Braz. J. Microbiol. 46(2), 465–476. http://doi.org/10.1590/s1517-838246220120018 (2015).

28. Cosate, M. R. V. et al. Molecular typing of Leptospira interrogans serovar Hardjo isolates from leptospirosis outbreaks in Brazilian livestock. BMC Vet. Res. 13(177), http://doi.org/10.1186/s12917-017-1081-9 (2017).

29. Zuerner, R. L. & Bolin, C.A. Differentiation of Leptospira interrogans isolates by IS1500 hybridization and PCR assays. J. Clin. Microbiol. 35(10), 2612–2617. http://doi.org/10.1128/jcm.35.10.2612-2617.1997 (1997).

30. Budihal, S.V. & Perwez, K. Leptospirosis Diagnosis: Competancy of Various Laboratory Tests. J. Clin. Diagn. Res. 8(1), 199–202. http://doi.org/10.7860/JCDR/2014/6593.3950 (2014).

31. La Peña-Moctezuma, A., Bulach, D. M., Adler, B. Genetic differences among the LPS biosynthetic loci of serovars of Leptospira interrogans and Leptospira borgpetersenii. Fems Immunol. Med. Mic. 31(1), 73–81. http://doi.org/10.1111/j.1574-695x.2001.tb01589.x (2001).

32. Stallman, N. D.. International Committee on Systematic Bacteriology Subcommittee on the Taxonomy of Leptospira: minutes of the meeting, 6 to 10 august 1982, boston, massachusetts. Int. J. Bacteriol. 34(2), 258–259. http://doi.org/10.1099/00207713-34-2-258 (1984).

33. Jolley, K.A. & Maiden, M.C.J. BIGSdb: Scalable analysis of bacterial genome variation at the population level. BMC Bioinformatics. 11(595), https://doi.org/10.1186/1471-2105-11-595 (2010).

34. Parra, G., Bradnam, K., Ning, Z., Keane, T., Korf, I. Assessing the gene space in draft genomes. Nucleic Acids Res. 37(1), 289–297. http://doi.org/10.1093/nar/gkn916 (2009).

35. Simão, F. A., Waterhouse, R. M., Ioannidis, P., Kriventseva, E. V., Zdobnov, E. M. BUSCO: assessing genome assembly and annotation completeness with single-copy orthologs. Bioinformatics. 31(19), 3210–3212. http://doi.org/10.1093/bioinformatics/btv351 (2015).

36. Emms, D. M. & Kelly, S. OrthoFinder: phylogenetic orthology inference for comparative genomics. Genome Biol. 20(238), https://doi.org/10.1186/s13059-019-1832-y (2019).

37. Fouts, D. E. The J. Craig Venter Institute (JCVI): lineardisplay. LinearDisplay. https://github.com/JCVenterInstitute/LinearDisplay (2019).

38. Altschul, S.F., Gish W., Miller W., Myers E.W., Lipman D.J. Basic local alignment search tool. J Mol Biol. 215(3), 403–10. https://doi.org/10.1016/S0022-2836(05)80360-2 (1990).

